# A phage-encoded counter-defense inhibits an NAD-degrading anti-phage defense system

**DOI:** 10.1101/2024.12.23.630042

**Authors:** Christian L. Loyo, Alan D. Grossman

## Abstract

Bacteria contain a diverse array of genes that provide defense against predation by phages. Anti-phage defense genes are frequently located on mobile genetic elements and spread through horizontal gene transfer. Despite the many anti-phage defense systems that have been identified, less is known about how phages overcome the defenses employed by bacteria. The integrative and conjugative element ICE*Bs1* in *Bacillus subtilis* contains a gene, *spbK*, that confers defense against the temperate phage SPβ through an abortive infection mechanism. Using genetic and biochemical analyses, we found that SpbK is an NADase that is activated by binding to the SPβ phage portal protein YonE. The presence of YonE stimulates NADase activity of the TIR domain of SpbK and causes cell death. We also found that the SPβ-like phage Φ3T has a counter-defense gene that prevents SpbK-mediated abortive infection and enables the phage to produce viable progeny, even in cells expressing *spbK*. We made SPβ-Φ3T hybrid phages that were resistant to SpbK-mediated defense and identified a single gene in Φ3T (*phi3T_120,* now called *nip* for NADase inhibitor from phage) that was both necessary and sufficient to block SpbK-mediated anti-phage defense. We found that Nip binds to the TIR (NADase) domain of SpbK and inhibits NADase activity. Our results provide insight into how phages overcome bacterial immunity by inhibiting enzymatic activity of an anti-phage defense protein.

**Author Summary:** Bacterial viruses (bacteriophages or phages) are widespread and abundant across the planet. Bacteria have a variety of immune systems, often found on mobile genetic elements, to combat phage predation. Phages can overcome these immune systems by mutating to avoid recognition or by producing molecules that prevent the immune system from working. We determined how an anti-phage defense system encoded by an integrative and conjugative element recognizes phage infection to cause cell death prior to the generation of phage progeny. We also identified a phage gene that prevents this defense system from functioning. The phage-encoded counter-defense protein inhibits the enzymatic activity of the anti-phage defense protein, enabling evasion of immunity and production of infectious phage. There are likely many different phage-encoded counter-defense genes yet to be discovered.

## Introduction

Bacteria have many different types of immune systems that protect against bacteriophages (phages). These anti-phage defense systems work through a variety of different mechanisms to limit the spread of phages. Many anti-phage defense systems work through abortive infection, wherein the phage and cell are both destroyed during the course of infection, preventing phage propagation within a population of cells [1]. Phages can have counter-defense genes that enable the phage to circumvent one or more anti-phage defense systems. These phage-encoded counter-defenses are diverse in how they enable evasion of the host-mediated defense, including blocking recognition of phage infection by the defense system and inhibiting the enzymatic activity of anti-phage defense proteins [2].

Bacteria typically contain several mobile genetic elements, including plasmids, temperate phages, and integrative and conjugative elements (ICEs). Anti-phage defense genes are often found on mobile genetic elements as part of the repertoire of so-called cargo genes. Cargo genes are not required for the lifecycle of a mobile element but often confer some benefit to their host. Because of their ability to move from host to host, mobile genetic elements that contain anti-phage defense genes facilitate rapid adaptation to phage infection and acquisition [3].

Many isolates of *Bacillus subtilis* harbor the integrative and conjugative element ICE*Bs1* [4]. ICE*Bs1* contains a gene, *spbK*, that causes abortive infection in response to infection by or activation of the temperate phage SPβ [5]. Here, we report that SpbK is an NADase that is activated by the phage portal protein YonE to cause NAD depletion during infection (**Fig 1A**). We also found that the SPβ-like phage Φ3T is resistant to SpbK-mediated abortive infection and identified a single gene in Φ3T, *phi3T_120* (now called *nip*, for NADase inhibitor from phage), that was both necessary and sufficient for this counter-defense. We found that Nip bound to the TIR (NADase) domain of SpbK and inhibited its NADase activity, preventing abortive infection and enabling the phage to grow and propagate (**Fig 1B**).

**Figure 1.**
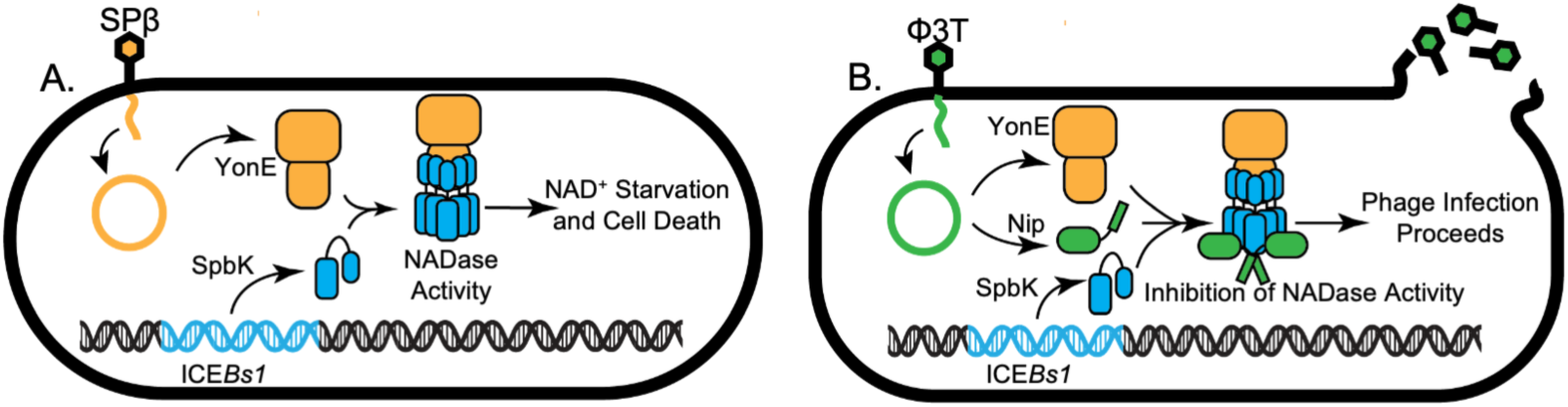
Model for activation and inhibition of the NADase SpbK. **A)** Model of SpbK-mediated anti-phage defense. SPβ phage adsorbs to the host cell and injects its DNA (orange) into the cell. Phage portal YonE (orange shapes) is produced during lytic phage infection. ICE*Bs1* (light blue) is an integrative and conjugative element found in the chromosome of *B. subtilis.* The ICE*Bs1*-encoded protein SpbK (light blue cylinders) associates with YonE and leading to activation of the SpbK NADase activity, NAD^+^ depletion, and cell death. **B)** Model of Nip-mediated counter-defense. The SPβ-like phage Φ3T (green) adsorbs to the cell and extrudes its DNA (green) into the cell. During lytic infection, Φ3T expresses YonE and a counter defense protein, Nip (green), that binds to SpbK (light blue) and inhibits SpbK-mediated NADase activity, resulting in phage production.

## Results

### The anti-phage defense protein SpbK is an NADase that is activated by the SPβ phage portal protein YonE

The product of the ICE*Bs1* gene *spbK* prevents propagation of the phage SPβ through an abortive infection mechanism that is activated by the phage protein YonE [5]. YonE is predicted to be the phage portal protein that is essential for DNA packaging into the capsid and subsequent extrusion of DNA from the capsid into the tail during infection [6,7]. SpbK contains a toll-interleukin-1 receptor-like (TIR) domain (**Fig 2A and 2B)** that is found in several proteins in both prokaryotes and eukaryotes [8,9], many of which have NADase activity [10].

**Figure 2.**
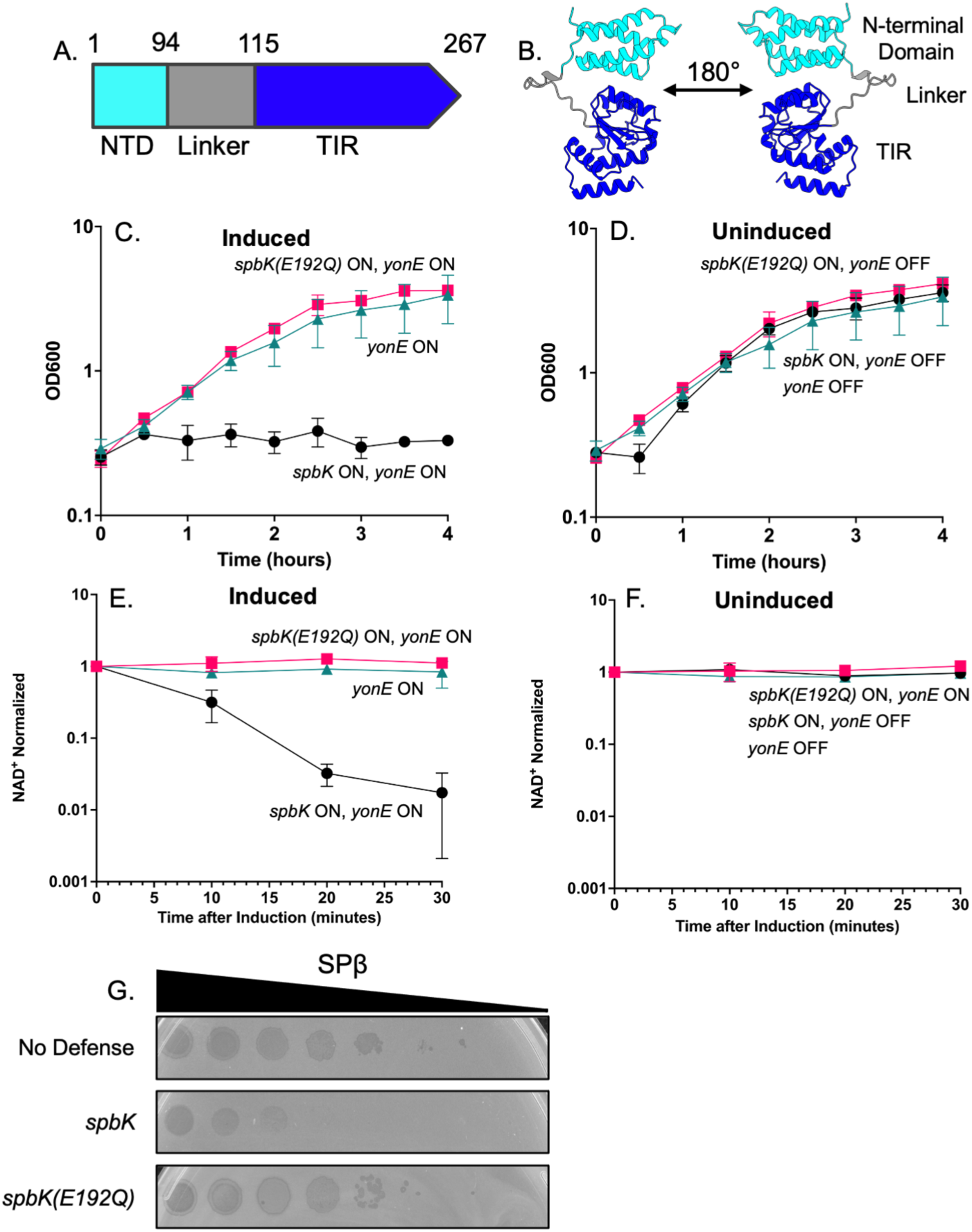
SpbK is an NADase that is activated by the phage protein YonE. **A)** Predicted domain architecture of SpbK. SpbK is 267 amino acids and includes the N-terminal domain (cyan; aa 1-94), the linker domain (aa 95-115), and the Toll-interleukin-1 receptor (TIR) domain (dark blue; aa 116-267). **B)** AlphaFold3 model of a monomer of SpbK. The N-terminal domain is colored in cyan, the linker domain is colored in gray, and the Toll-interleukin-1 receptor (TIR) domain in dark blue. The predicted template modeling score (pTM) is 0.62 and the mean predicted local distance test (pLDDT) score is 95.53, indicating high confidence in the predictions. **C, D, E, F**) Strains co-expressing *spbK* and *yonE* (CMJ685, black circles), *spbK*(*E192Q*) and *yonE* (CLL317, pink squares), or only *yonE* (CMJ616, turquoise triangles) were grown in S7_50_ minimal medium. Expression of Pspank(hy)-*yonE* was induced with 1 mM IPTG. Culture turbidity (**C, D**) and levels of NAD^+^ (**E, F**) were measured over time. Levels of NAD^+^ were normalized to the amount of NAD^+^ per OD 0.025 cells at T=0. Measurements at T=0 were taken immediately before addition of IPTG. Error bars represent standard deviation and are not always visible. **G**) Ten-fold serial dilutions of SPβ phage were spotted onto isogenic strains expressing no anti-phage defense (CU1050, top row), *spbK* (CMJ534, middle row), or *spbK*(*E192Q*) (CLL164, bottom row). Large zones of clearing are indicative of a confluence of phage plaques and cell lysis. Small zones of clearing are indicative of individual or small clusters of phage plaques.

We found that SpbK is an NADase that is activated by YonE. We measured NAD^+^ levels in cells following production of SpbK and YonE separately and together. We expressed *spbK* from its endogenous promoter (PspbK*-spbK*) and *yonE* from the IPTG-inducible promoter Pspank(hy) (Pspank(hy)*-yonE*). Expression of *spbK* or *yonE* alone had no detectable effect on growth (**Fig 2C and 2D**), as previously determined [5], and no detectable effect on cellular levels of NAD^+^ (**Fig 2E and 2F**). In marked contrast, expression of both *spbK* and *yonE* together caused growth arrest (**Fig 2C and 2D**), as previously described [5], and a drop in levels of NAD^+^ (**Fig 2E and 2F**). By 15 minutes after inducing expression of *yonE* by addition of IPTG, there was an approximate 10-fold drop in levels of NAD^+^ and by 30 minutes, there was an approximately 100-fold drop in levels of NAD^+^ (**Fig 2E and 2F**).

TIR-domain containing proteins that act as NADases have a conserved glutamate in the active site that is required for cleavage of NAD^+^ [9–11]. A multiple sequence alignment of SpbK to similar proteins indicated that a highly conserved glutamate at position 192 (Glu192) of SpbK was likely to be in the catalytic site. We changed Glu192 to glutamine (E192Q) and tested for effects on cell growth (**Fig 2C and 2D**), levels of NAD^+^ (**Fig 2E and 2F**), and anti-phage defense (**Fig 2G**). The *spbK*(*E192Q*) mutation abolished anti-phage defense (**Fig 2G**). That is, SPβ was able to grow and make plaques on cells expressing the mutant *spbK*, in contrast to the absence of phage growth in cells expressing wild type *spbK* (**Fig 2G**). In contrast to the results with co-expression of *yonE* with wild type *spbK* (**Fig 2C**), co-expression of *yonE* with *spbK*(*E192Q*) caused no detectable change in cell growth (**Fig 2C**) and no decrease in the amount of NAD^+^ (**Fig 2E**), indicating that both of these phenotypes are dependent on the activity of the TIR domain in SpbK. Together, these results indicate that SpbK acts as an NADase and is activated by the SPβ phage portal protein YonE.

### SpbK and YonE interact directly

We tested for interaction between SpbK and YonE using immunoprecipitation of epitope-tagged proteins. We fused a FLAG tag to the N-terminus of SpbK (FLAG-SpbK) and a 3xMyc tag to the C-terminus of YonE (YonE-Myc). Both fusion proteins were functional (**S1A Fig**): FLAG-SpbK inhibited growth of SPβ (**S1A Fig**) and expression of YonE-Myc caused growth arrest in cells also expressing SpbK (**S1B Fig**).

We found that SpbK and YonE interact directly *in vivo*. We co-expressed FLAG-SpbK and YonE-Myc in the absence of any other genes from ICE*Bs1* or SPβ, immunoprecipitated FLAG-SpbK with anti-FLAG antibodies, ran the samples on SDS polyacrylamide gels, and probed Western blots for YonE-Myc using anti-c-Myc antibodies. We found that YonE-Myc co-precipitated with FLAG-SpbK (**Fig 3A, lane 4**). In control experiments in which we expressed SpbK (no tag) with YonE-Myc or FLAG-SpbK with YonE (no tag), we no longer detected YonE-myc in Western blots of immunoprecipitated material (**Fig 3A, lanes 2 and 3**). These controls indicate that the anti-FLAG antibodies were specifically immunoprecipitating FLAG-SpbK and the anti-Myc antibodies were specifically detecting YonE-Myc. Further, the presence of YonE-Myc in the IP fraction was dependent on FLAG-SpbK, as we detected YonE-Myc in input samples (whole cell lysates) but only detected YonE-Myc in the immunoprecipitates when FLAG-SpbK was also present (**Fig 3A, lanes 2 and 3**). Together, these results indicate that SpbK and YonE are likely interacting directly, and that this interaction does not require any other ICE*Bs1* or SPβ products.

**Figure 3.**
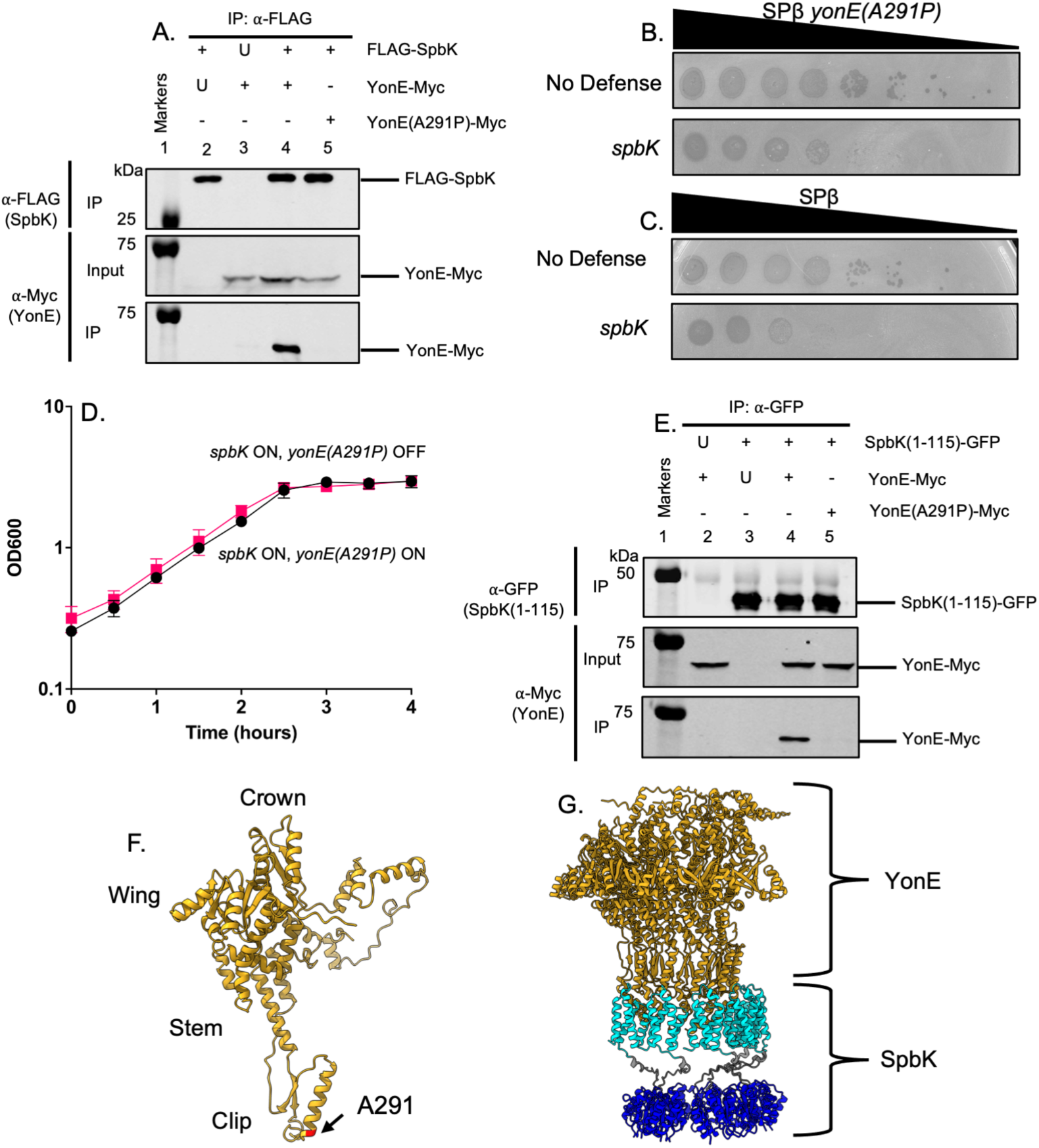
The N-terminal domain of SpbK interacts with the clip domain of YonE. In the immunoprecipitation experiments (**A, E**), the indicated proteins were produced in cells without ICE*Bs1* or SPβ. **A**) SpbK interacts with YonE. Co-immunoprecipitation experiments to probe interactions between FLAG-SpbK and YonE-Myc or YonE(A291P)-Myc. Top row: Western blot probed with α-FLAG monoclonal antibody on IP fractions. Second row: Western blot probed with α-c-Myc monoclonal antibody on input fractions. Third row: Western blot probed with α-c-Myc monoclonal antibody on IP fractions. Lane 1 contains molecular weight markers from the Odyssey One-Color Protein Molecular Weight Marker Ladder (LI-COR). Lysates from the following strains were used for immunoprecipitations: CLL609 (lane 2), CLL334 (lane 3), CLL373 (lane 4), and CLL615 (lane 5). Data shown are representative of three biological replicates. U, untagged protein expressed; +, tagged protein expressed; -, protein not expressed. **B, C**) Ten-fold dilutions of SPβ *yonE*(*A291P*) (**B**) or wild type SPβ (**C**) were spotted onto isogenic strains with no phage defense (CU1050, top row) or expressing *spbK* (CMJ534, bottom row). Large zones of clearing are indicative of a confluence of phage plaques and cell lysis. Small zones of clearing are indicative of individual or small clusters of phage plaques. **D)** Expression of *yonE*(*A291P*) does not activate SpbK-mediated growth arrest. Strains co-expressing *spbK* and *yonE*(*A291P*) were grown in S7_50_ minimal medium and growth was monitored by OD_600_ for 4 hours. *yonE*(*A291P*) was induced with 1mM IPTG (black circles) or left uninduced (pink squares). Measurements at T=0 were taken immediately before addition of IPTG. Data shown are from three biological replicates. Error bars represent standard deviation and are not always depicted due to the size of the data point. **E)** The N-terminal domain of SpbK interacts with YonE, but not YonE(A291P). Co-immunoprecipitation experiments to probe interactions between SpbK(1-115)-GFP and YonE-Myc or YonE(A291P)-Myc. Top row: Western blot probed with α-GFP polyclonal antibody on IP fractions. Second row: Western blot probed with α-c-Myc monoclonal antibody on input fractions. Third row: Western blot probed with α-c-Myc monoclonal antibody on IP fractions. Lane 1 contains molecular weight markers from the Odyssey One-Color Protein Molecular Weight Marker Ladder (LI-COR). Lysates from the following strains were used for immunoprecipitations: CLL718 (lane 2), CLL719 (lane 3), CLL720 (lane 4), and CLL721 (lane 5). Data shown are representative of three biological replicates. U, untagged protein expressed; +, tagged protein expressed; -, protein not expressed. **F)** Alanine 291 is located in the clip domain of YonE. A monomer of the YonE portal protein was modeled using AlphaFold3. The approximate location of the clip, stem, wing, and crown domains are labeled. The location of the A291P mutation is highlighted in red. The pTM score is 0.75 and the mean pLDDT score is 77.91, indicating moderate confidence in the predictions. **G)** The N-terminal domain of SpbK is predicted to interact with the clip domain of YonE. Six units of YonE (gold) were modeled with six units of SpbK using AlphaFold3. The N-terminal region of SpbK is colored cyan, the linker region of SpbK is colored gray, and the TIR domain of SpbK is colored dark blue. The pTM score is 0.47, the mean pLDDT score is 67.77, and the interface predicted template modeling (ipTM) score is 0.43, indicating low confidence in the specific molecular interactions.

Because YonE from SPβ activates SpbK to cause abortive infection preventing phage growth [5], we sought to isolate phage mutants that do not activate SpbK. Such mutants should be able to grow and make plaques on cells producing SpbK. We isolated such a mutant from a population of approximately 10^10^ phage (**Fig 3B)**. This phage had a mutation in *yonE* that changed alanine at position 291 to proline {*yonE*(*A291P*)}. Relative to that of the wild type phage, the SPβ *yonE*(*A291P*) mutant had increased plaquing efficiency on cells expressing *spbK*, although it did not fully escape SpbK-mediated defense (**Fig 3C)**. Expression of the mutant *yonE* allele did not cause growth arrest when co-expressed with *spbK* (**Fig 3D)**, indicating that YonE(A291P) might not interact with or has a weaker interaction with SpbK than wild type YonE.

We used immunoprecipitation to test for interaction between SpbK and the YonE(A291P) mutant. As above, we expressed FLAG-SpbK and YonE(A291P)-Myc and immunoprecipitated FLAG-SpbK. We did not detect YonE(A291P)-Myc in the immunoprecipitates as determined by Western blot (**Fig 3A, lane 5**). Together, these results indicate that SpbK interacts with YonE, and that the YonE(A291P) mutant that is functional for phage production has reduced interaction with SpbK.

We found that an N-terminal fragment of SpbK was sufficient to interact with YonE. We fused the N-terminal 115 amino acids of SpbK, including the N-terminal domain and linker region (**Fig 2A**) but not the TIR domain, to the N-terminus of GFP (SpbK(1-115)-GFP) and expressed this allele in cells with YonE-Myc. We immunoprecipitated with anti-GFP antibodies and found that YonE-Myc co-precipitated (**Fig 3E, lane 4**), indicating this region of SpbK directly interacts with YonE. As controls, we expressed SpbK(1-115)-GFP with wild type YonE (no tag) **(Fig 3E, lane 2)** and the N-terminal region of SpbK without a GFP tag with YonE-Myc **(Fig 3E, lane 3)**. As above, we detected YonE-Myc in input samples, and detection of YonE-Myc in the immunoprecipitates was dependent on expression of SpbK(1-115)-GFP. Similar to results expressing FLAG-SpbK and YonE(A291P)-Myc, we did not observe an interaction between SpbK(1-115)-GFP and YonE(A291P)-Myc (**Fig 3E, lane 5)**. Because the TIR domain was no longer present, co-expression of SpbK(1-115) or SpbK(1-115)-GFP did not cause growth arrest. Further, SpbK(1-115) and SpbK(1-115)-GFP was not sufficient to inhibit production of SPβ following infection (**S1A Fig**), indicating that this interaction alone was not sufficient to block phage production.

We compared AlphaFold3 [12] predictions to our experimental observations of interaction between YonE and SpbK (**Fig 3F and 3G**). The YonE(A291P) mutation is predicted with high confidence to be in the clip domain (**Fig 3F**). The clip domain of portal proteins interacts with the large terminase and plug proteins during DNA packaging and helps stabilize the phage tail [13], and YonE-like portal proteins form dodecamers [7]. Due to the current limitations of the AlphaFold3 server, we were unable to model the complete dodecameric structure with an equivalent number of SpbK monomers. Therefore, we modeled six YonE monomers with six SpbK monomers. AlphaFold3 predicted, with low confidence, that the N-terminal region of SpbK interacts with the clip domain of YonE, with the interface located near A291 (**Fig 3G**). The low confidence of the model indicates that the molecular details of the known interaction are likely not accurate, perhaps due to limitations on the oligomeric inputs. Nonetheless, the experimental results (**Fig 3A, E**) are quite clear about interactions between the N-terminal domain of SpbK and YonE and the role of the clip domain in those interactions. Oligomerization of TIR domains is often a prerequisite for NADase activity [14–16], and interaction between SpbK and YonE likely facilitates interactions between the TIR domains to activate the NADase. Overall, our results are most consistent with the conclusion that the N-terminal region of SpbK interacts with the clip domain of YonE, and this interaction leads to oligomerization and activation of SpbK, and subsequent depletion of NAD (**Fig 1A**).

### Phage Φ3T encodes a counter-defense against SpbK

We tested the ability of SpbK to protect against the phages Φ3T and ρ11, both closely related to SPβ [17,18]. Both Φ3T and ρ11 were able to grow and make plaques with similar efficiencies on strains with and without *spbK* (**Fig 4A and S2AB**), indicating that both phages were resistant to SpbK-mediated defense. It was possible that resistance of these two phages to SpbK was caused by a mutation in *yonE*, analogous to the mutant we isolated above. However, we found that the sequence of *yonE* from each phage was identical at the nucleotide sequence level to that of *yonE* from SPβ. Thus, we suspected that Φ3T and ρ11 might encode a counter-defense that prevented SpbK from functioning in abortive infection.

**Figure 4.**
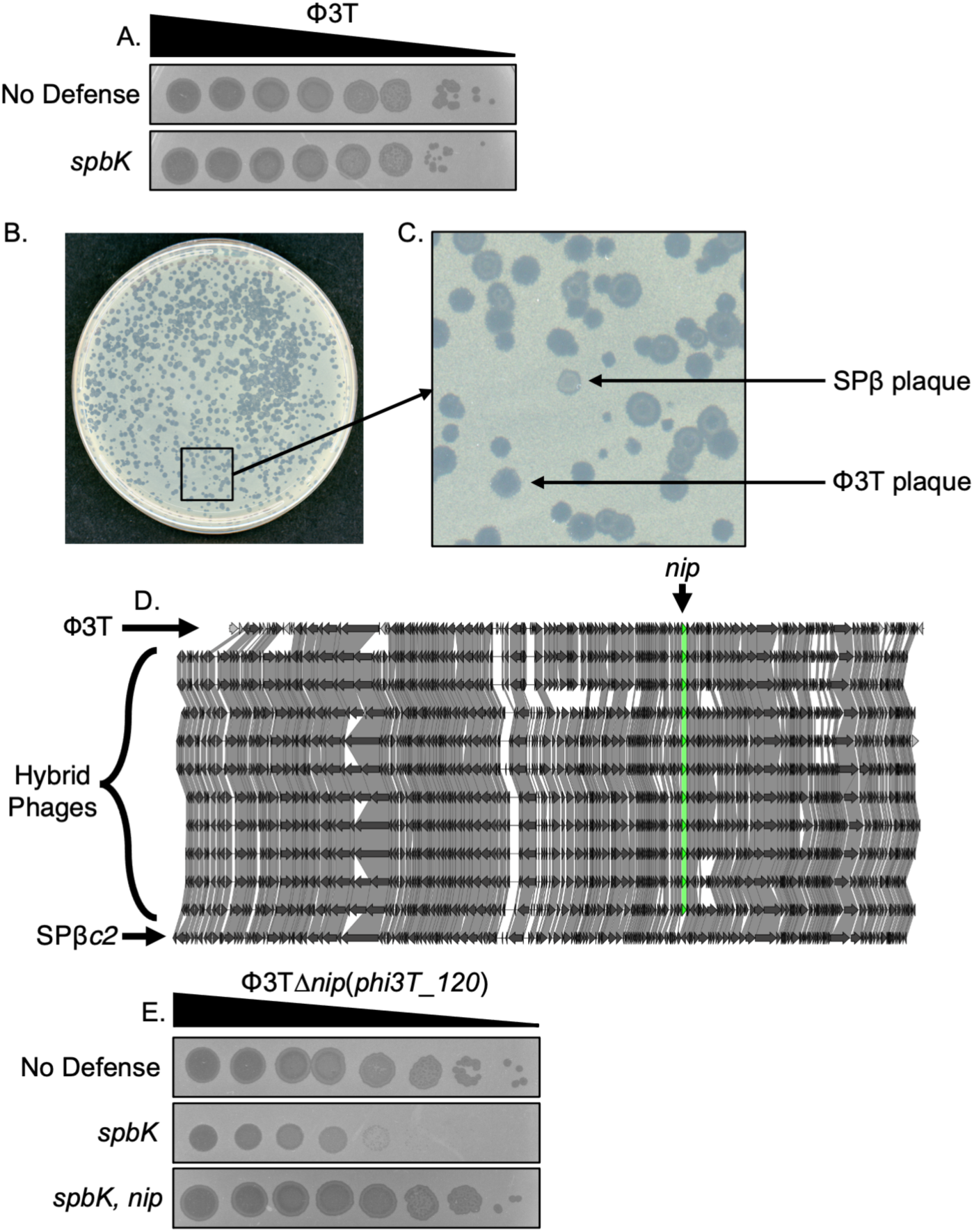
Phage Φ3T encodes a counter-defense against SpbK-mediated phage defense. **A)** Φ3T escapes SpbK-mediated phage defense. Ten-fold dilutions of Φ3T was spotted onto isogenic strains without anti-phage defense (CU1050; top row) or expressing *spbK* (CMJ534; bottom row). Large zones of clearing are indicative of a confluence of phage plaques and cell lysis. Small zones of clearing are indicative of individual or small clusters of phage plaques. **B)** Plaques formed on a plate with strain expressing *spbK*. Double lysogens of Φ3T and SPβ*c2* were treated with heat shock (methods). Phages produced from strain CLL182 were mixed with strain CMJ534 (*spbK*+) and plated via top agar overlay. **C)** Magnified view of an area from (**B**) containing different plaque morphologies. SPβ-like plaques (small and cloudy) could be distinguished from Φ3T plaques (large and clear) visually. Arrows indicate typical plaque morphologies of SPβ and Φ3T. **D)** Genome comparisons of Φ3T (top), Φβ phages (Φ3T-SPβ hybrids) (middle), and SPβ*c2* (bottom). Genomes of hybrid phage lysogens were sequenced and annotated. Prophage sequences were manually extracted from the genome and aligned with clinker [45]. The counter-defense gene *nip* (*phi3T_120*) is highlighted in green. **E)** Nip is necessary and sufficient for evading SpbK-mediated phage defense. Ten-fold dilutions of Φ3T (left) and Φβ3 (Φ3T-SPβ hybrid phage) (right) were spotted onto isogenic strains without anti-phage defense (CU1050; top row), expressing *spbK* (CMJ534; middle row), or co-expressing *spbK* and *nip* (CLL356; bottom row). Large zones of clearing are indicative of a confluence of phage plaques and cell lysis. Small zones of clearing are indicative of individual or small clusters of phage plaques.

#### Φ3T-SPβ hybrid phages

To identify the possible counter-defense gene(s), we made hybrid phages, each of which had some genes from SPβ and some from Φ3T. To do this, we isolated SPβ-like hybrids that were able to evade SpbK-mediated defense. Briefly, we constructed a double lysogen that harbored both Φ3T and a temperature sensitive mutant of SPβ, SPβ*c2* [19]. Since ρ11 contains the same attachment site as SPβ at *spsM* [17,20] and Φ3T integrates at *kamA* [17], we focused on Φ3T since we could easily construct a double lysogen. From the double lysogen, we induced the temperature sensitive SPβ*c2* with heat shock. The population of phages coming from the double lysogen should include SPβ*c2* and Φ3T, and some recombinants between the two phages. We screened this mixed population of phages for those that had the small, turbid plaque morphology of SPβ (in contrast to the much larger plaques formed by Φ3T) but were able to grow and make plaques on cells expressing *spbK* (**Fig 4B and 4C**). We isolated 10 independent hybrid phages (named Φβ1 - Φβ10), verified the phenotypes, and then sequenced and compared their genomes (**Fig 4D**).

There were two genes from Φ3T, *phi3T_120* and *phi3T_121* that were present in all 10 hybrid phages (**Fig 4D**). We changed the third codon of *phi3T_121* from a lysine to a stop codon{Φ3T *phi3T_121*(*K3\**). We found that this mutant was virtually indistinguishable from wild type Φ3T in its resistance to SpbK-mediated defense **(S2C and S2D Fig)**, indicating that *phi3T_121* was not needed for counter-defense. We therefore focused on *phi3T_120*.

*phi3T_120* (*nip*) is required for counter-defense.

We found that *phi3T_120* (*nip*) was required for counter-defense to SpbK. Deletion *phi3T_120* (Φ3T Δ*phi3T_120*) caused reduced plaquing efficiency on cells expressing *spbK* (**Fig 4E and S2B**), indicating that *phi3T_120* (called hereafter as *nip*, for NADase inhibitor from phage) was required for Φ3T to overcome SpbK-mediated defense and productively grow on cells expressing *spbK*. ρ11 has a gene that is identical to *nip* (*phi3T_120*) from Φ3T. We deleted *nip* from ρ11 (ρ11 *Δnip*) and found that this mutant phage had decreased efficiency of plating on cells expressing *spbK* (**S2A and S2B Fig**), indicating that *nip* also provides ρ11 with counter-defense against SpbK.

#### *phi3T_120* (*nip*) is sufficient for counter-defense

We found that expression of *nip* was sufficient to enable SPβ to form plaques on cells expressing *spbK*. We cloned *nip* under control of the constitutive promoter Ppen (Ppen-*nip*) and integrated this in the *B. subtilis* chromosome. SPß was able to grow and make plaques on cells that were expressing both *spbK* and *nip* (**S2B Fig**), in contrast to the reduced plaquing efficiency on cells expressing only *spbK* (**Fig 2G and S2B**). Based on these results, we conclude that *nip* is a counter-defense gene that enables resistance to SpbK-mediated defense.

Together, these results demonstrate that the Φ3T gene *nip* (*phi3T_120*) is required for Φ3T to evade SpbK-mediated defense and is the only gene in Φ3T needed to confer this counter-defense to SPβ.

### Nip inhibits the NADase activity of SpbK

We found that *nip* prevented growth arrest and NAD^+^ depletion caused by co-expression of *spbK* and *yonE*. We expressed *nip* (Ppen-*nip*) in the absence of any phage genes, other than *yonE*, in cells expressing *spbK* and *yonE* (**Fig 5A**). The growth of cells expressing all three genes was indistinguishable from that of cells expressing none or each gene individually (**Fig 5A**; **1C and 1D**). Additionally, there was no detectable drop in levels of NAD^+^ in cells expressing all three genes (*nip*, *spbK*, and *yonE*) (**Fig 5B**). These results are in marked contrast to the effects of expressing *spbK* and *yonE* in the absence of *nip* (**Fig 2C**) and indicate that *nip* was functional outside the context of phage infection and was sufficient to inhibit the NADase activity of SpbK and to confer counter-defense.

**Figure 5.**
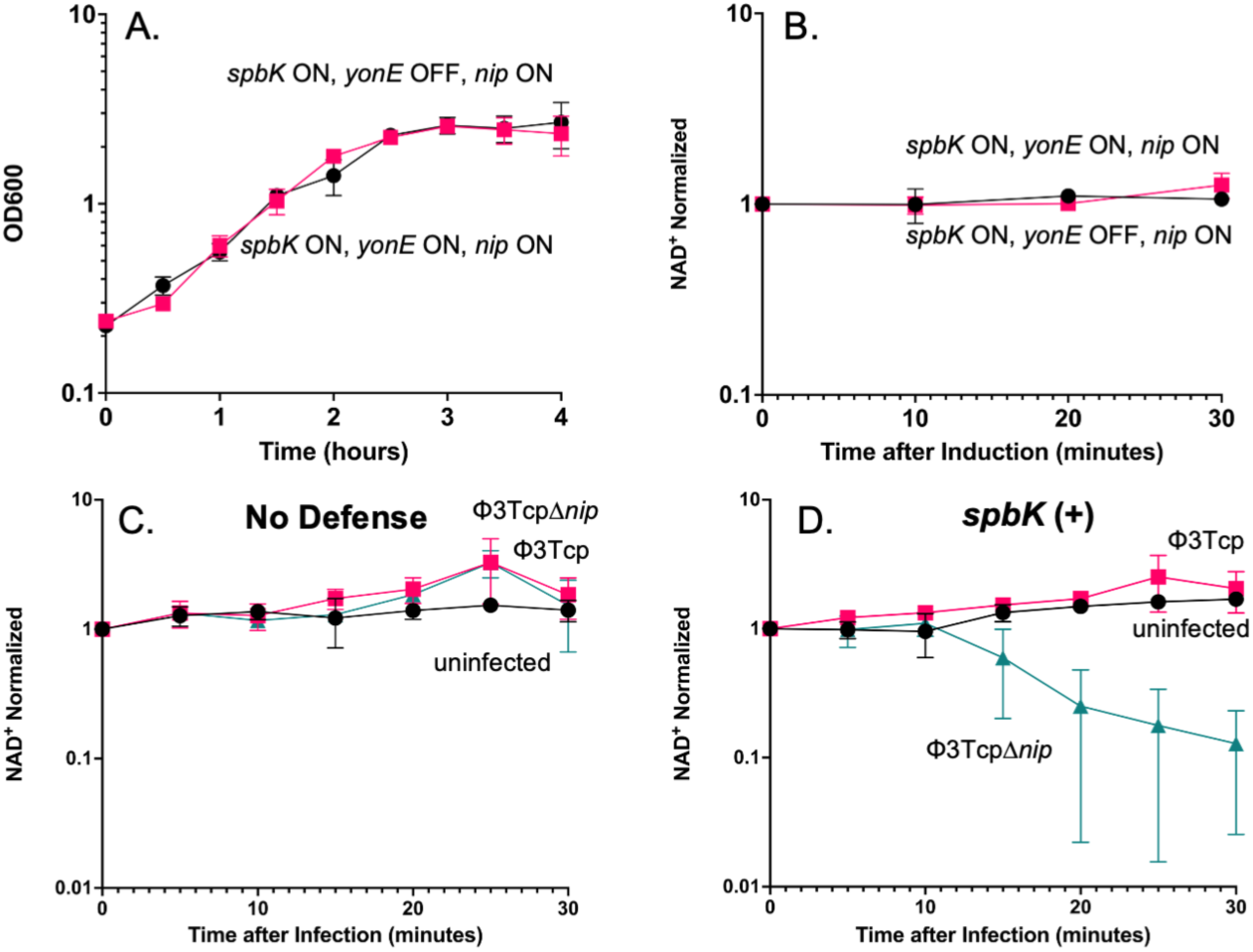
Expression of *nip* prevents growth arrest and NAD^+^ depletion during phage infection. **A, B**) Expression of *nip* prevents growth arrest (**A**) and depletion of intracellular levels of NAD^+^ (**B**) when strains are also expressing *spbK* and *yonE*. Culture turbidity by OD600 (**A**) and intracellular levels of NAD^+^ (**B**) were measured and followed over time in strains co-expressing *spbK, yonE,* and *nip* (CLL353; black circles) or strains co-expressing *spbK* and *nip*, with uninduced *yonE* (CLL353; pink squares) in S7_50_ minimal medium. Expression of *yonE* was induced by addition of 1mM IPTG. Data represent three biological replicates. Error bars represent standard deviation and are not always depicted due to the size of the data point. **C, D**) *nip* prevents depletion of NAD^+^ in cells during phage infection. Strains without phage defense (PY79) (**C**) or expressing *spbK* (CMJ684) (**D**) were infected with Φ3T*cp* (pink squares), Φ3T*cp* Δ*nip* (turquoise triangles), or left uninfected (black circles). Levels of NAD^+^ were measured and followed over time. Samples at T=0 were taken immediately before infection. Phages were added at a multiplicity of infection of 10 in infected samples. Data shown represent three biological replicates. Error bars represent standard deviation and are not always depicted due to the size of the data point.

We also found that *nip* functions to inhibit the NADase activity of SpbK during phage infection. As described above, infection of *spbK*-expressing cells with Φ3T resulted in phage growth and eventual cell lysis, leading to the release of functional phage. We found no detectable drop in levels of NAD^+^ when cells expressing *spbK* were infected with a clear plaque mutant of Φ3T (Φ3Tcp) which can only undergo lytic growth and cannot enter the lysogenic cycle (**Fig 5D**). In contrast, there was an approximate 10-fold drop in levels of NAD^+^ when *spbK*-expressing cells were infected by Φ3Tcp Δ*nip* (**Fig 5D**). Based on these results, we conclude that *nip* is a counter-defense gene in Φ3T (and ρ11) that is both necessary and sufficient for inhibiting the NADase activity of SpbK and enabling productive phage growth in cells with the SpbK abortive infection system.

### Nip binds to SpbK and forms a tripartite complex with YonE

There are a few different ways in which Nip could directly affect the NADase activity of SpbK. Nip might interact directly with YonE and prevent it from interacting with SpbK. Nip might instead interact with SpbK and prevent it from interacting with YonE. In this model, Nip would likely interact with the N-terminal domain of SpbK (the region that interacts with YonE). Alternatively, Nip might interact with the C-terminal TIR domain of SpbK and inhibit NADase activity. This interaction might or might not prevent interactions between SpbK and YonE.

To distinguish between these models, we used epitope-tagged proteins (described below) to test directly for interactions between Nip, SpbK, and YonE, in the absence of any other phage or ICE*Bs1* genes. Experiments described below demonstrate that Nip interacts with the C-terminal domain of SpbK and that YonE, SpbK, and Nip can form a tripartite complex.

#### Direct interaction between Nip and SpbK

We found that Nip interacts with SpbK. We fused a 3xMyc tag to the C-terminus of Nip (Nip-Myc). Nip-Myc was functional in preventing SpbK-mediated defense against SPβ (**S1A Fig**). We immunoprecipitated FLAG-SpbK using anti-FLAG antibodies from cells co-expressing Nip-Myc and found that that Nip-Myc co-immunoprecipitated (**Fig 6A, lane 4**), indicating that Nip and SpbK interact. In control experiments in which we expressed wild type SpbK (no tag) with Nip-Myc or FLAG-SpbK with wild type Nip (no tag) we no longer detected Nip-Myc in Western blots of immunoprecipitated material (**Fig 6A, lanes 2 and 3**), indicating the antibodies used in the immunoprecipitation and Western blot were specific to the tagged proteins and not cross-reacting with the other protein of interest.

**Figure 6.**
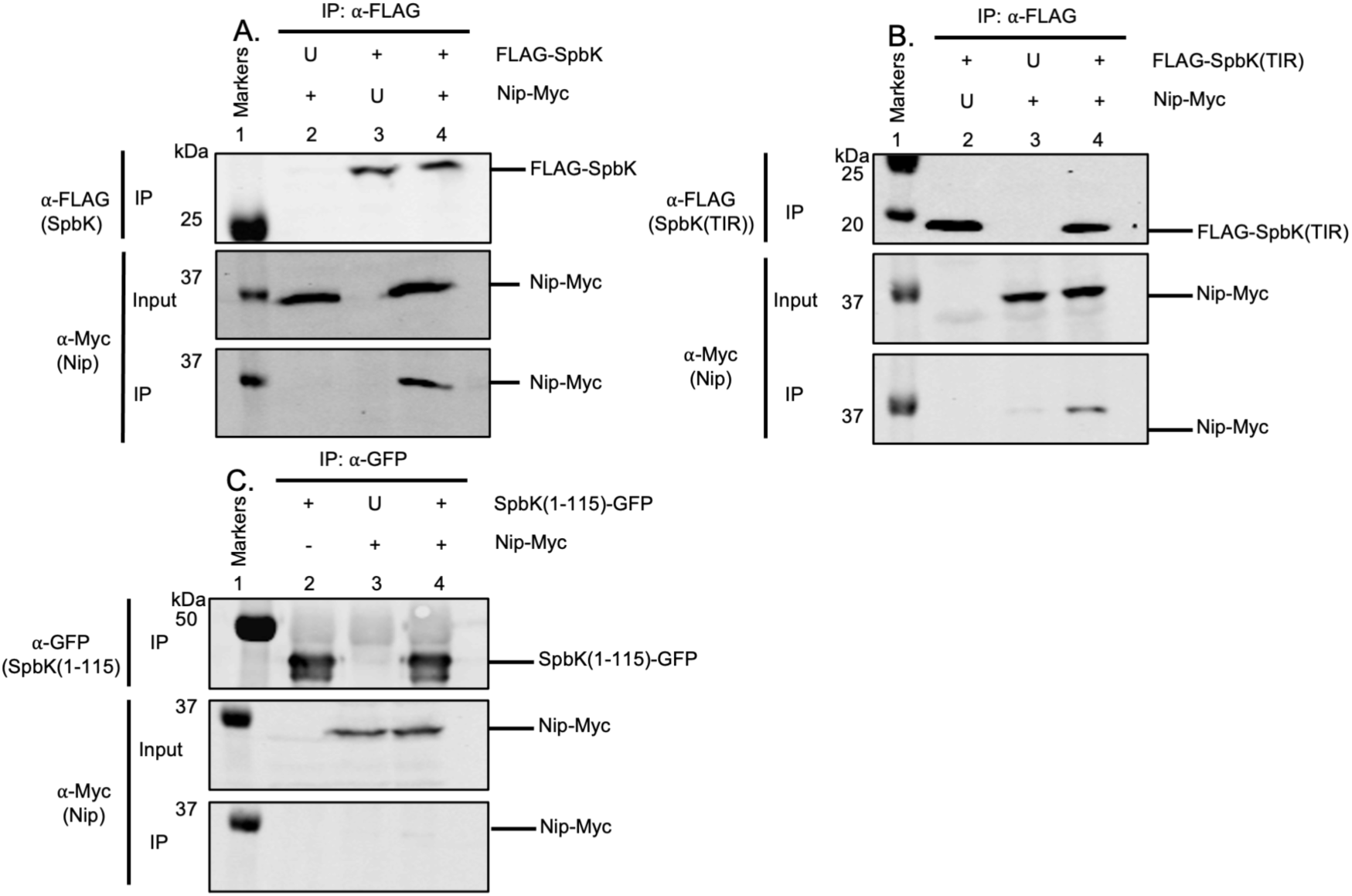
Nip interacts with the TIR domain of SpbK. Indicated proteins were produced in cells without ICE*Bs1* or SPβ. In each panel, lane 1 contains molecular weight markers from the Odyssey One-Color Protein Molecular Weight Marker Ladder (LI-COR). U, untagged protein expressed; +, tagged protein expressed; -, protein not expressed. **A)** SpbK interacts with Nip. Monoclonal α-FLAG antibodies were used to immunoprecipitate FLAG-SpbK. Western blots were probed with α-FLAG monoclonal antibodies (top panel) or α-c-Myc monoclonal antibodies (bottom two panels). Top row: Western blot probed with α-FLAG monoclonal antibody on immunoprecipitated fractions. Second row: Western blot probed with α-c-Myc monoclonal antibody on input fractions. Third row: Western blot probed with α-c-Myc monoclonal antibody on IP fractions. Lysates from the following strains were used for immunoprecipitations: CLL388 (lane 2), CLL390 (lane 3), and CLL376 lane 4). Data shown are representative of three biological replicates. **(B)** Nip interacts with the TIR domain of SpbK. Monoclonal α-FLAG antibodies were used to immunoprecipitated FLAG-SpbK. Top row: Western blot probed with α-FLAG monoclonal antibody on immunoprecipitated fractions. Second row: Western blot probed with α-c-Myc monoclonal antibody on input fractions. Third row: Western blot probed with α-c-Myc monoclonal antibody on immunoprecipitated fractions. Lysates from the following strains were used for immunoprecipitations: CLL669 (lane 2), CLL671 (lane 3), and CLL670 (lane 4). Data shown are representative of three biological replicates. (C) No interaction was detected between Nip and the N-terminal domain of SpbK. Polyclonal α-GFP antibodies were used to immunoprecipitate SpbK(1-115)-GFP. Top row: Western blot probed with α-GFP polyclonal antibody on immunoprecipitated fractions. Second row: Western blot probed with α-c-Myc monoclonal antibody on input fractions. Third row: Western blot probed with α-c-Myc monoclonal antibody on immunoprecipitated fractions. Lysates from the following strains were used for immunoprecipitations: CLL726 (lane 2), CLL725 (lane 3), and CLL727 (lane 4). Data shown are representative of three biological replicates.

#### Nip interacts with the C-terminal NADase/TIR domain of SpbK

We found that Nip interacts with the C-terminal part of SpbK that contains the TIR domain. We fused the FLAG tag to the N-terminus of the linker and TIR domain (aa 95-276) of SpbK {FLAG-SpbK(TIR)} and precipitated with anti-FLAG antibodies, as with full length FLAG-SpbK above. We found that Nip-Myc co-immunoprecipitated with FLAG-SpbK(TIR) (**Fig 6B, lane 4**). As controls, we expressed FLAG-SpbK(TIR) with wild type Nip (no tag) (**Fig 6B, lane 2**) or SpbK(TIR) (no tag) with Nip-Myc (**Fig 6B, lane 3**) and did not observe an interaction. Nip-Myc was detected in in cell extracts (**Fig 6B, lanes 3 and 4**) and only detected in immunoprecipitates when FLAG-SpbK(TIR) was also present (**Fig 6B, lane 4**).

We also tested for the ability of Nip to interact with the N-terminal region of SpbK. We immunoprecipitated the SpbK(1-115)-GFP fusion protein with anti-GFP antibodies and did not detect coprecipitation of Nip in cells also expressing Nip-Myc (**Fig 6C, lane 4**). Similar to controls in previous experiments, we expressed SpbK(115)-GFP with wild type Nip (no tag) (**Fig 6C, lane 2**) and SpbK(1-115) (no tag) with Nip-Myc (**Fig 6C, lane 3**) and did not observe an interaction. Together, the results from the immunoprecipitation experiments indicate that Nip interacts with the TIR domain and not the N-terminal region of SpbK.

#### Nip forms a tripartite complex with SpbK and YonE

We found that Nip, SpbK, and YonE coprecipitate (**Fig 7, S3 Fig**). Using cells expressing Nip-GFP, FLAG-SpbK, and YonE-Myc, we immunoprecipitated Nip-GFP and found that both FLAG-SpbK and YonE-Myc were in the immunoprecipitates (**Fig 7, lane 5**). The presence of FLAG-SpbK and YonE-Myc was specific as we detected little or none of either of these proteins in the immunoprecipitates from cells expressing Nip without the GFP tag (**Fig 7, lane 3**). Both proteins were present in the cell extracts prior to immunoprecipitation (**S3 Fig**). Additionally, the antibodies used in the Western blots were specific for each of the tagged proteins as there was no signal if the protein did not contain the epitope tag (**Fig 7, lane 2, no FLAG; lane 4, no Myc**).

**Figure 7.**
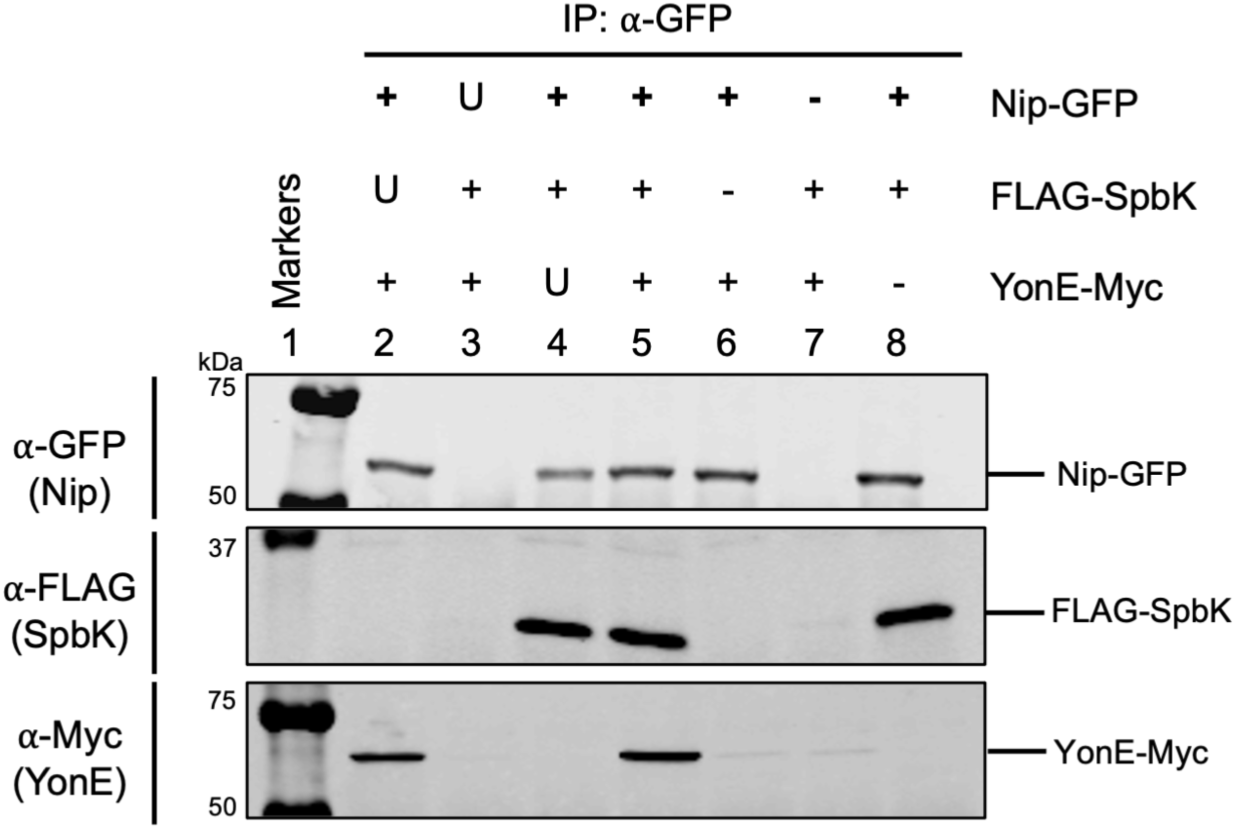
Nip, SpbK, and YonE form a tripartite complex. Indicated proteins were produced in cells without ICE*Bs1* or SPβ. α-GFP antibodies were used to immunoprecipitate Nip-GFP. Top row: Western blot probed with α-GFP polyclonal antibody on immunoprecipitated fractions. Second row: Western blot probed with α-FLAG monoclonal antibody on immunoprecipitated fractions. Third row: Western blot probed with α-c-Myc monoclonal antibody on immunoprecipitated fractions. The ladder in lane 1 is composed of molecular weight markers from the Odyssey One-Color Protein Molecular Weight Marker Ladder (LI-COR). Lysates from the following strains were used for immunoprecipitations: CLL633 (lane 2), CLL642 (lane 3), and CLL704 (lane 4), CLL498 (lane 5), CLL497 (lane 6), CLL373 (lane 7), and CLL382 (lane 8). Data shown are representative of three biological replicates. U, untagged protein expressed; +, tagged protein expressed; -, protein not expressed.

Co-precipitation of Spbk and YonE with Nip could indicate separate bipartite interactions: Nip with SpbK, as described above (**Fig 6**), and Nip with YonE. If so, then coprecipitation of YonE with Nip would be independent of the presence of SpbK, as the coprecipitation of Nip and SpbK is independent of YonE (**Fig 6**). Alternatively, the coprecipitation of YonE with Nip could reflect the presence of a tripartite complex containing YonE, SpbK, and Nip. In this scenario, coprecipitation of YonE with Nip would depend on the presence of SpbK.

We found that Nip, SpbK, and YonE form a tripartite complex. YonE-Myc was not coprecipitated with Nip-GFP in cells that were not also producing SpbK (Fig 7, lane 6). The presence of either FLAG-SpbK (**Fig 7 lane 5**) or the native untagged SpbK (**Fig 7 lane 2**) was required for coprecipitation of YonE-Myc with Nip-GFP. Together, these results indicate that Nip, SpbK, and YonE can form a tripartite complex (**Fig 7**), that interactions between Nip and SpbK are independent of YonE (**Fig 6**), and interactions between SpbK and YonE are independent of Nip (**Fig 3**).

In total, the simplest interpretation of our results is that the clip domain of the phage portal protein YonE interacts with the N-terminal domain of the ICE*Bs1*-encoded protein SpbK. This interaction stimulates the NADase activity of SpbK, leading to a drop in intracellular levels of NAD^+^, cell death, and the inability of phage to grow and make plaques (**Fig 1A**). Phages that encode the counter-defense protein Nip (or when Nip is ectopically expressed) escape the SpbK-mediated defense because Nip binds to the TIR domain of SpbK and inhibits NADase activity and abortive infection (**Fig 1B**).

## Discussion

Work presented here shows that the anti-phage defense protein SpbK, encoded by ICE*Bs1*, is an NADase that is activated by the essential phage portal protein YonE, encoded by the termperate phage SPβ and its relatives, including phages Φ3T and ρ11. We identified a counter-defense gene, *nip*, found in Φ3T and ρ11. Nip interacts directly with the TIR domain of SpbK to inhibit NADase activity. Below we discuss the presence of anti-phage defense genes on mobile genetic elements, the types of phage products that are often involved in activating defense systems, the role of NAD^+^ in anti-phage defense, and the roles of phage-encoded counter-defense genes.

### Anti-phage defense genes on mobile genetic elements

Anti-phage defense systems are often found on mobile genetic elements [21,22]. Mobile genetic elements such as ICE*Bs1* facilitate rapid acquisition and dissemination of accessory genes, such as anti-phage defense systems. Turnover of anti-phage defense systems has been observed to occur rapidly and is mediated by the movement of mobile genetic elements in *Vibrio* [3,23]. The spread of mobile genetic elements throughout a population of bacteria likely enables population-based anti-phage defense strategies like abortive infection to be more effective in preventing phage outbreaks.

Anti-phage defense genes on mobile genetic elements have been observed to be located in recombinational hotspot regions. P2 and P4-like phages in *E. coli* contain several anti-phage systems across diverse isolates [24]. The location of anti-phage hotspots between core conserved genes likely facilitates recombination between similar elements, increasing genetic diversity within bacterial populations. Overall, we anticipate many undiscovered anti-phage systems to be found in mobile genetic elements across different bacteria.

### Phage products that activate anti-phage defense systems

Anti-phage systems that work through abortive infection are inactive until phage infection is recognized. The triggers for abortive infection are often conserved phage-encoded proteins that are essential for the phage lifecycle. For example, the phage portal protein (YonE in the case of SPβ) is essential for phage growth. Most mutations in an essential phage gene would prevent phage growth (beneficial for the bacterium), which makes it a suitable target for detecting phage infection. The low frequency (∼10^-10^) with which we found the SPβ *yonE*(*A291P*) mutant that escapes SpbK-mediated defense is likely due to the fitness tradeoffs imposed on the phage when both the function as a phage portal and escaping anti-phage defense are required of YonE. The frequency of phages acquiring new genes is likely greater than the relatively low frequency of mutations that allow escape from defense and maintain full function for phage growth. This is likely the reason that some phages in the SPβ family have a counter-defense gene rather than an alteration in *yonE* to prevent SpbK-mediated abortive infection.

### Role of NAD^+^ in anti-phage defense

Anti-phage defense systems that halt cell growth and/or cause cell death target essential host functions. NAD^+^ is required in all known living organisms and is the target of several anti-phage defense systems. For example, the Thoeris anti-phage system has a TIR-domain-containing protein, ThsB, that generates a signaling molecule that activates ThsA, a protein with a SIR2 domain, that then depletes NAD^+^ in infected cells [9,25]. Some of the cyclic-oligonucleotide-based anti-phage signaling systems (CBASS) also contain TIR domains with NADase activity [26,27]. Some prokaryotic Argonaute systems contain TIR domains that cause depletion of NAD^+^ when foreign DNA is present in cells [28]. TIR domain-containing proteins are widespread and play important roles in regulating cell death in both prokaryotes and eukaryotes [8].

### Phage-encoded counter-defense systems

Several different phage-encoded counter-defense systems counteract the effects of anti-phage systems that have NADases. For example, Tad1 and Tad2 inhibit Thoeris-mediated anti-phage defense by sequestering the signaling molecules produced by Thoeris during infection, thus preventing activation of the Thoeris-encoded NADase [2,29]. Another strategy is to replenish NAD^+^ levels during phage infection after an NADase effector from an anti-phage defense has been activated. The NARP1 and NARP2 counter-defense systems work in this way [30]. Notably, the NARP1 system is encoded by an SPβ-like phage that does not contain *nip*. Thus, related phages have two different mechanisms of counteracting NADase mediated defense: replenish NAD levels (NARP1) and directly bind and inhibit the NADase (Nip).

*nip* is in a region of Φ3T that is expressed early (based on analyses of SPβ), in contrast to *yonE* which is expressed late during phage infection [31]. Expression of *nip* before *yonE* likely provides sufficient time for Nip to bind and inactivate the TIR (NADase) domain of SpbK before YonE is expressed. The operon containing *nip* encodes another counter-defense, Gad1, which inhibits Gabija anti-phage defense [2]. Other phages have counter-defense genes that are genetically linked. For example, phage T4 genes *tifA* and *dmd* are in an operon and each confers resistance to toxin-antitoxin systems [32,33]. Similarly to anti-phage defense hotspots, it appears that there are hotspots in phages for the accommodation of counter-defense genes, and these are often located in operons that are expressed early during phage infection, likely to inhibit anti-phage systems prior to or concomitant with expression of the activator protein.

## Materials and methods

### Media and growth conditions

*B. subtilis* cells were grown in LB or S7_50_ minimal medium supplemented with 0.1% glutamate and 1% glucose as a carbon source [34]. *E. coli* was grown in LB medium and on LB agar plates (1.5% bacto-agar). Isopropyl β-D-1-thiogalactopyranoside (IPTG) was used at 1 mM to induce expression from the LacI-repressible IPTG-inducible promoter Pspank(hy). Antibiotics used for selection in *B. subtilis* included: kanamycin (5 µg/ml), spectinomycin (100 µg/ml), chloramphenicol (5 µg/ml), tetracycline (10 µg/ml), and erythromycin (0.5 µg/ml) plus lincomycin (12.5 µg/ml) for macrolide-lincosamide-streptogramin (MLS) resistance.

For typical experiments, a culture was started from a single colony (after overnight growth) and grown at 37℃ to mid-late exponential phase in the indicated medium. Cells were diluted into fresh medium for continued growth as indicated. For experiments with genes expressed from an inducible promoter, cultures were split at the time of dilution into medium with and without inducer, as indicated.

Temperate phages were induced by addition of mitomycin C (MMC) to 1 µg/ml to a lysogen growing exponentially in LB medium at 37℃. After MMC induction, cells were grown for one additional hour and pelleted at 4000g for 5 minutes in a tabletop centrifuge. The lysate was transferred to a new tube, 1:100 v/v chloroform was added to inhibit cell growth, and stored at 4℃.

#### Efficiency of plating

Phage stocks were diluted in phage buffer (150 mM NaCl, 40 mM Tris-Cl, 10 mM MgSO_4_) and mixed with 300 µl of cells that were growing exponentially in LB medium. Cells and phage were incubated for 5 minutes at room temperature to allow adsorption, transferred to 3 ml of molten LB + 0.5% bacto-agar, and plated onto warm LB agar plates. Plates were incubated at 30℃ overnight to allow plaque formation.

Serial dilutions (typically 10-fold) for spot assays were made in phage buffer and 2 µl were spotted onto a lawn of cells on LB agar plates.

### Cell growth following phage infection

Cells were grown from a single colony in LB medium to OD_600_ ∼0.5, and back diluted to OD ∼0.05, and 180 µl were added to each well of a 96 well plate and mixed with 20 µl of phage or 20 µl of phage buffer. Plates were incubated at 37℃ with shaking in a Biotek Synergy H1 plate reader with OD_600_ measurements taken every 15 minutes.

### Strains and alleles

*E. coli* strain AG1111 (MC1061 F’ *lacI*^q^ *lacZM15* Tn*10*) was used as a host for plasmid cloning. *B. subtilis* strains used are listed in Table 1. All strains are derivatives of PY79 [35] or CU1050 [36], both of which are cured of SPβ and ICE*Bs1*. Indicated alleles were introduced by natural transformation [37] and appropriate selection. New constructs and alleles are described below.

**Table 1.**
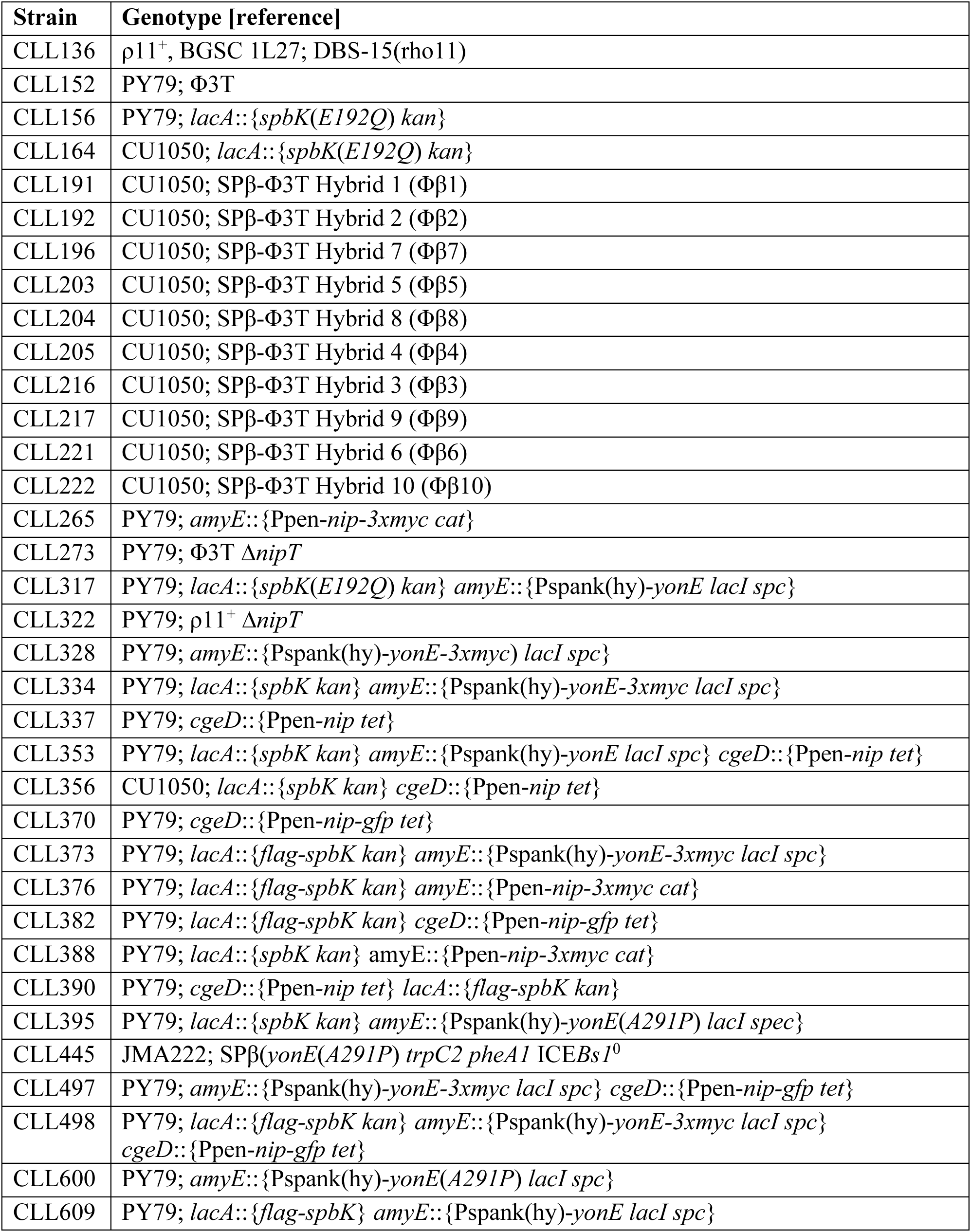
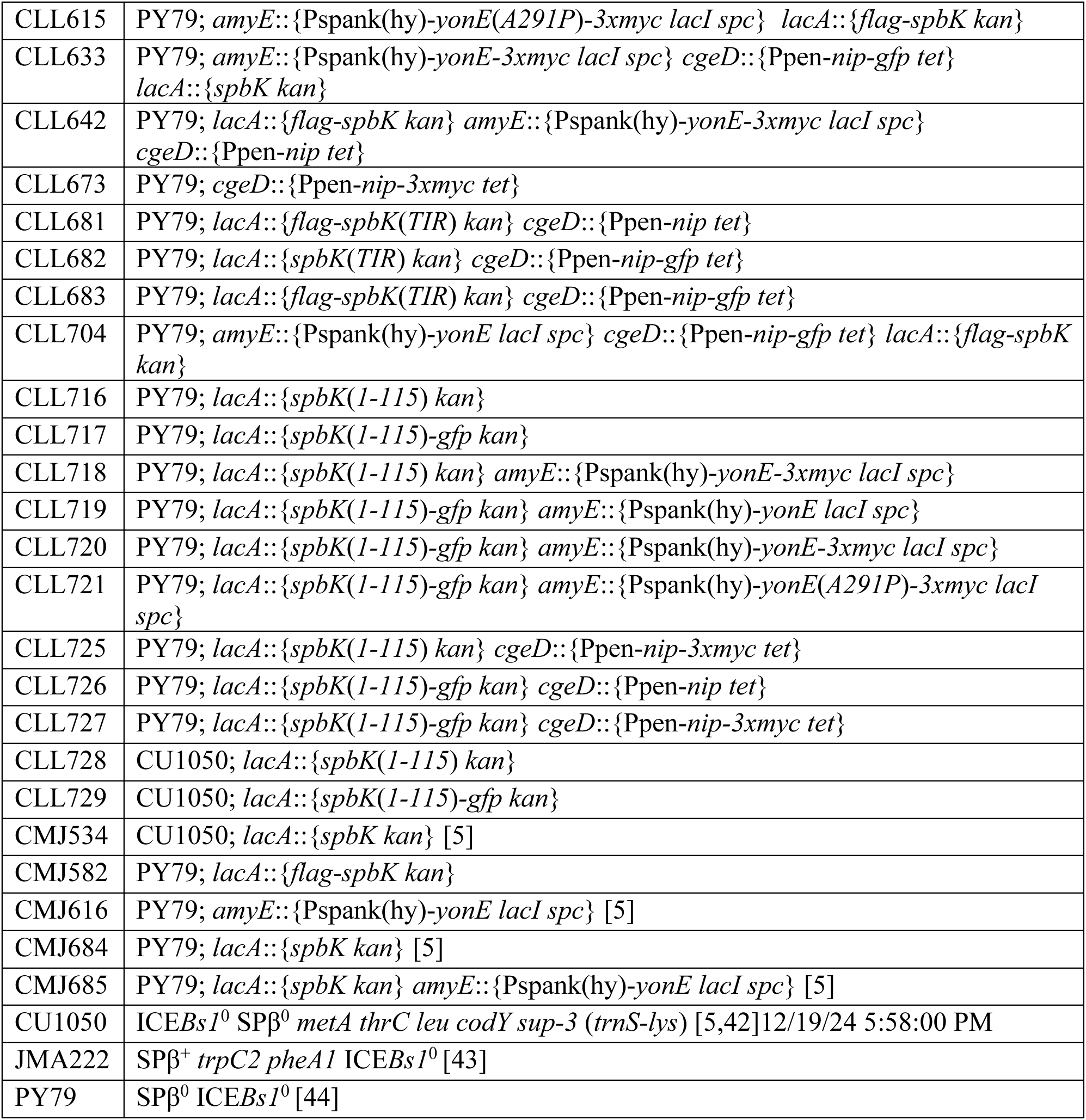
*B. subtilis* strains used.

**Table 2.** Phages used.

| Phage | Genotype, phenotype, description (comment, source, reference) |
| --- | --- |
| SPβ | from strain JMA222 (Auchtung et al., 2005) |
| SPβ*c2* | ts mutant of SPβ; temperature-inducible; (phage isolated from BGSC 1A322; CU4142) |
| SPβ*cp* | Spontaneous clear plaque mutant of SPβ; (this study) |
| SPβ *yonE(A291P)* | SPβ mutant that escapes SpbK-mediated defense; in strain CLL445 (this study) |
| Φ3T | from strain CLL152; {phage isolated from BGSC 1L26; 3610(phi3T)} |
| Φ3T*cp* | Spontaneous clear plaque mutant of Φ3T (this study) |
| Φ3TΔ*nip* | From strain CLL273 (this study) |
| Φ3T*cp*Δ*nip* | Spontaneous clear plaque mutant of Φ3TΔ*nip*; (this study) |
| ρ11 | from strain CLL136; {phage isolated from BGSC 1L27; DBS-15(rho11)} |
| ρ11Δ*nip* | from strain CLL322 (this study) |
| $\Phi\beta$1 | SPβ-Φ3T hybrid; from strain CLL191 (this study) |
| $\Phi\beta$2 | SPβ-Φ3T hybrid; from strain CLL192 (this study) |
| $\Phi\beta$3 | SPβ-Φ3T hybrid; from strain CLL216 (this study) |
| $\Phi\beta$4 | SPβ-Φ3T hybrid; from strain CLL205 (this study) |
| $\Phi\beta$5 | SPβ-Φ3T hybrid; from strain CLL203 (this study) |
| $\Phi\beta$6 | SPβ-Φ3T hybrid; from strain CLL221 (this study) |
| $\Phi\beta$7 | SPβ-Φ3T hybrid; from strain CLL196 (this study) |
| $\Phi\beta$8 | SPβ-Φ3T hybrid; from strain CLL204 (this study) |
| $\Phi\beta$9 | SPβ-Φ3T hybrid; from strain CLL217 (this study) |
| $\Phi\beta$10 | SPβ-Φ3T hybrid; from strain CLL222 (this study) |

Unmarked deletions were generated by inserting flanking homology regions into pCAL1422 digested with BamHI and EcoRI via Gibson isothermal assembly [38]. pCAL1422 is a plasmid containing *E. coli lacZ* and *cat* (chloramphenicol-resistance in *B. subtilis*) [39]. The assembled plasmids were transformed into *E. coli* strain AG1111. After the correct plasmid construct was verified by PCR and Sanger sequencing, plasmids were transformed into *B. subtilis* selecting for chloramphenicol resistance (indicating an integrated plasmid by single crossover recombination). Transformants were grown in LB liquid medium and plated onto LB plates supplemented with 120 mg/ml X-gal. Colonies were screened for loss of *lacZ* and verified to have the desired deletion by PCR.

#### Deletion of *nip* from Φ3T

We made a deletion of the Φ3T gene *nip* in a lysogen of Φ3T. The deletion of *nip* extended from the start to stop codons of *nip* and was constructed as described above after cloning DNA flanking the deletion endpoints into the integration vector pCAL1422.

#### Ectopic expression of *nip*

*cgeD*::Ppen*-nip* (CLL337) was used to test for inhibition of SpbK-mediated anti-phage defense and growth arrest. *nip* was cloned downstream of the promoter Ppen and the *spoVG* ribosome binding site (Ppen*-nip*) and integrated into the nonessential gene *cgeD.* The 5’ fragment was amplified containing the upstream *cgeD* homology arm, tetracycline resistance gene, Ppen, and the *spoVG* ribosome binding site. The 3’ fragment contained the downstream *cgeD* homology arm. *nip* from 3 bp upstream of the start codon to 110 bp downstream of the stop codon was amplified by PCR, assembled with the 5’ and 3’ fragments by Gibson isothermal assembly, and transformed into *B. subtilis*.

#### Fusions of *nip* to *3xmyc* and *gfp*

Constructs expressing *nip* with different epitope tags were used in immunoprecipitation assays. Briefly, gDNA from CLL337 was used to amplify fragments containing *nip*, Ppen promoter, *spoVG* ribosome binding site, and cgeD or amyE homology arms. DNA encoding the tags was amplified from previously generated constructs and inserted at the 3’ end of *nip* via Gibson isothermal assembly to generate *nip-3xmyc* (CLL265 and CLL673) and *nip-gfp* (CLL370) [38]. The resulting constructs were transformed into *B. subtilis* selecting for tetracycline resistance.

#### Ectopic expression alleles of *spbK*

Strains expressing *spbK* that had been integrated into *lacA* were previously described [5]. *lacA*::{*spbK*(*E192Q*) *kan*} (CLL156), encoding a catalytically dead mutant of SpbK, was made by amplifying two fragments of *spbK* and surrounding *lacA* sequence (with genomic DNA from CMJ684 [5] as the template), upstream and downstream of the mutation, using overlapping primers that contained the mutation (changing codon 192 from 5’-GAA-3’ to 5’-CAA-3’). Fragments were assembled via isothermal assembly and the resulting construct was transformed into *B. subtilis,* selecting for resistance to kanamycin. kanamycin. *lacA*::{*flag-spbK kan*} (CMJ582), used in immunoprecipitation assays, was constructed by Chris Johnson. It consists of a *flag* tag inserted directly after the start codon of *spbK*.

#### Ectopic expression alleles of *yonE*

*amyE*::{Pspank(hy)*-yonE spc*} (CMJ616) was previously described [5]. *amyE*::{Pspank(hy)-*yonE*(*A291P*)} (CLL380) was made by amplifying a fragment of *yonE*(*A291P*) from the mutant SPβ that was resistant to (escaped) SpbK-mediated defense and inserted into pCAL1422 via Gibson isothermal assembly, essentially as described generally above. The plasmid was crossed into a strain with amyE::{Pspank(hy)*-yonE* spc} (CMJ616) and recombinants that had lost the plasmid were screened by PCR to have the desired *yonE*(*A291P*) mutation.

*amyE*::{Pspank(hy)*-yonE-3xmyc spc*} (CLL328) and *amyE*::{Pspank(hy)*-yonE*(*A291P*)*-3xmyc spc*} (CLL600) were used for co-immunoprecipitation assays. For each allele, a *3xmyc* tag was fused to the 3’ end of *yonE*, directly upstream of the stop codon.

#### Ectopic expression of the N-terminal region of SpbK

*lacA*::{*spbK*(*1-115*) *kan*} (CLL716) was used as a control in immunoprecipitation assays. The 5’ end of *spbK* was amplified by PCR. This fragment extends from 330 bp upstream of the start codon to codon 115 and includes the *spbK* promoter and encodes the N-terminal and linker regions of SpbK.

*lacA*::{*spbK*(*1-115*)*-gfp kan*} (CLL717) was used in immunoprecipitation assays. Fragments were amplified from CLL716 and *gfp* inserted between codon 115 and the stop codon of *spbK*.

#### Ectopic expression of the TIR domain of SpbK

*lacA*::{*spbK*(*TIR*) *kan*} (CLL662) and *lacA*::{*flag-spbK*(*TIR*)} (CLL661) were used in immunoprecipitation assays. The linker and TIR domains (encoded by codons 95-267) were expressed under the endogenous promoter of *spbK*. Fragments were amplified from CMJ582 (*lacA*::{*flag-spbK kan*}) extending from the 5’ *lacA* homology arm to the *flag tag* for the first fragment and extending from codon 95 to the 3’ *lacA* homology arm for the second fragment. This construct was transformed into *B. subtilis*, selecting for kanamycin resistance.

#### Isolation of clear plaque phage mutants

All phages used in this study are temperate and typically develop turbid plaques on susceptible cells. Occasionally, clear plaques are observed, indicative of a mutant that can only undergo lytic growth and cannot enter the lysogenic lifestyle. Clear plaque mutant phages were isolated by picking a spontaneously occurring clear plaque, resuspended in phage buffer, and used to infect cells to allow phage growth. Infected cells were added to top agar (LB + 0.5% agar), plated via double agar overlay method, and incubated overnight. Phage buffer (5 ml) was added to the plates and the cells within top agar were scraped off. The mix was centrifuged and chloroform added to the supernatant (phage stock) to inhibit cell growth. Phage stocks were stored at 4℃.

#### Lysogens of temperate phages

Cells were plated with phage in top agar (LB + 0.5% agar) to generate turbid phage plaques (indicative of lysogens). Plaques were picked with a toothpick and streaked onto LB plates to isolate single colonies. Strains were confirmed to have the desired phage via PCR.

### Generation and sequencing of hybrid phages

Hybrid phages were generated inducing SPβ*c2*-Φ3T double lysogens with heat shock at 50℃ for 20 minutes. Cells were grown for an additional hour at 37℃ to allow phage production. Phage lysate containing approximately 10^3^ PFUs was mixed with 3ml of molten top agar (LB + 0.5% agar), poured onto LB plates, and incubated overnight at 30℃ overnight to allow the formation of plaques. Lysogens were isolated from SPβ-like plaques (Fig 4C) and genomic DNA was prepared using the DNeasy Tissue and Blood Kit (Qiagen) and sequenced via Nanopore sequencing on an R9 PromethION flowcell. Genomes were assembled using Flye assembler [40] and annotated using Prokka [41].

### Co-immunoprecipitation and Western blotting

Cells were grown in 100 ml LB cultures to OD 0.5-1.0 and 50 OD units of cells were pelleted in conical tubes at 4000*g* for 10 minutes, washed once with 5 ml of PBS, spun once more at 4000*g* for 10 minutes, and the supernatant was decanted. The pellet was resuspended in 450 µl of resuspension buffer (20% sucrose, 50 mM NaCl, 10 mM EDTA, 10 mM Tris-HCl pH 8.0) supplemented with 10 mg/ml lysozyme, 10 µl of P8849 protease inhibitor cocktail, and 250 units of Benzonase and incubated at 37℃ for 15 minutes. 450 µl of lysis buffer (100 mM Tris-HCl pH 7.0, 300 mM NaCl, 10 mM EDTA, 1% Triton X-100) was used to complete lysis. 30 µl of protein A beads suspended in ddH_2_O (Fisher Scientific, 50% v/v) were used to clear the lysate for non-specific binding proteins. Beads were pelleted, supernatant transferred to a new tube and incubated with appropriate antibody for 30 minutes. Antibodies used to bind the protein A beads include rabbit anti-GFP (Covance) and rabbit anti-FLAG (Cell Signaling Technologies). After incubation with antibody, 60 µl of protein beads suspended in ddH_2_O (Fisher Scientific, 50% v/v) were added and incubated overnight at 4℃ to form the immunocomplex. Beads were then washed three times with wash buffer (22.5mM Tris-HCl (pH 8.0), 0.5M Sucrose, 10mM EDTA) supplemented with 1% triton. 2x SDS sample buffer (100 mM Tris-HCl (pH 6.8), 20% glycerol, 4% SDS, 0.02% bromophenol blue) supplemented with 2-mercaptoethanol was added and boiled to elute proteins from the beads. Proteins were separated on a 10% SDS-PAGE and transferred to 0.2 µm nitrocellulose membrane using the Trans-blot SD semi-dry transfer cell (Bio-Rad). Blots were blocked for one hour with Odyssey Blocking Buffer. Mouse M2 anti-flag (1:1000; Millipore Sigma), rabbit anti-GFP (1:10,000; Covance), or mouse anti-c-Myc (1:500; Thermo Fischer) antibodies were used as primary antibodies. Blots were then washed with PBST (phosphate buffered saline + 0.2% Tween) five times for five minutes. Goat anti-mouse antibodies with a conjugated fluorophore (1:10,000; Avantor/VWR) was used as a secondary antibody. Blots were imaged on a Li-Cor scanner.

### Measuring NAD^+^ levels

NAD^+^ was measured using the NAD/NADH-Glo Assay kit (Promega). Cells were grown overnight at 30°C in S7_50_ minimal medium supplemented with glucose. Starter cultures were grown in S7_50_ supplemented with glucose at 37℃ until OD_600_ 0.5 and diluted into S7_50_ supplemented with glucose to an OD_600_ of 0.25. Where indicated, regulated promoters were induced by addition of 1mM IPTG and 5 ml samples were taken every 10 minutes after induction and chilled on ice to stop growth. Cells were pelleted for 5 minutes at 10000*g* at room temperature on a tabletop centrifuge, supernatant was decanted, and cells were saved at −80℃ until processing. Cells were processed following the supplier’s protocol.

During phage infection, cells were infected at a multiplicity of infection of 10, harvested every 5 minutes, immediately chilled on ice to stop growth, and spun at 4000*g* for 5 minutes at room temperature. Cells were processed as above.

## Acknowledgements

We thank Stuart Levine and the MIT BioMicro Center for help with genome sequencing, the Bacillus Genetic Stock Center for providing strains, and members of the Grossman Lab for some strains and helpful discussions and Michael Laub, Gene-Wei Li, and Andrew Camilli for useful discussions.

---

**S1 Fig.**
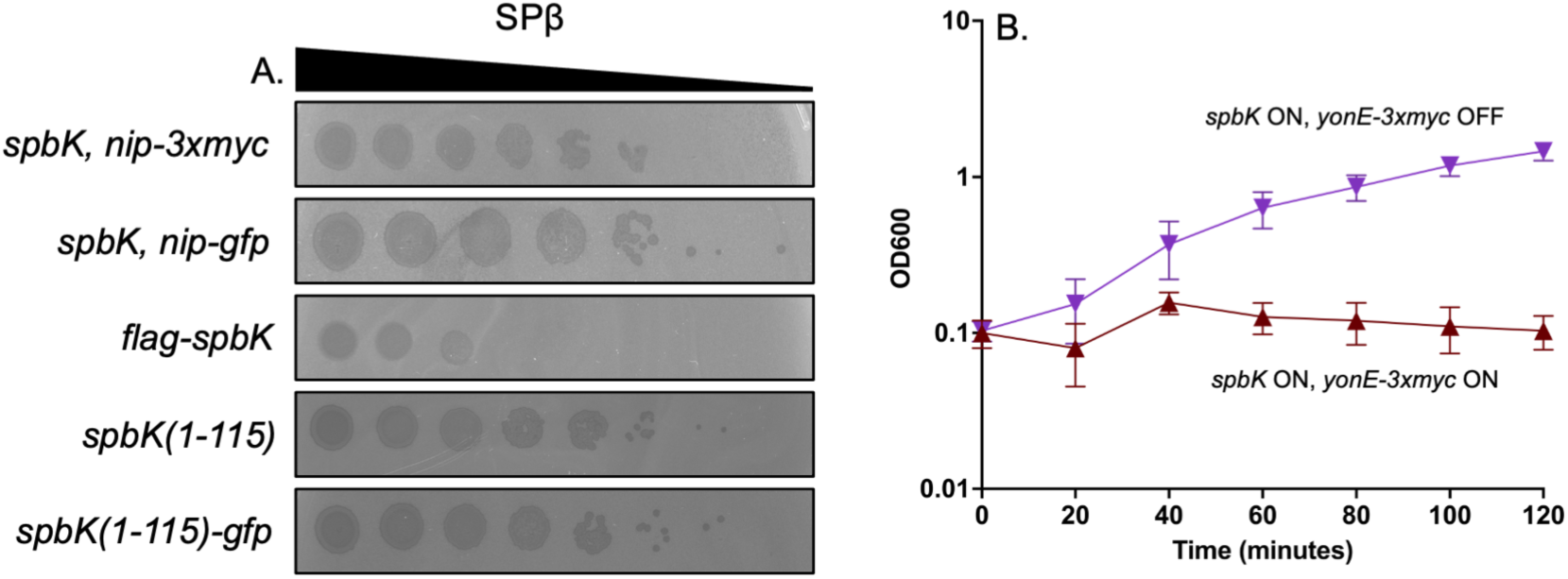
Fusion proteins of Nip, SpbK, and YonE are functional. **A)** Fusion proteins Nip-Myc (top row) and Nip-GFP (second row) were co-expressed with wild type SpbK. FLAG-SpbK (third row), SpbK(1-115) (fourth row), and SpbK(1-115)-GFP (fifth row) were expressed alone. Ten-fold dilutions of SPβ were spotted and functionality was assessed by observing presence or absence of phage plaques. Large zones of clearing are indicative of a confluence of phage plaques and cell lysis. Small zones of clearing are indicative of individual or small clusters of phage plaques. **B)** YonE-Myc causes growth arrest when co-expressed with wild type SpbK. Strains co-expressing *spbK* and *yonE-3xmyc* were grown in LB medium. Turbidity of the culture was measured by OD_600_ and followed over time when *yonE-3xmyc* was induced with 1mM IPTG (maroon triangles) or left uninduced (purple inverted triangles). Measurements at T=0 were taken immediately before addition of IPTG. Error bars represent standard deviation. Data shown are from three biological replicates. Error bars represent standard deviation and are not always depicted due to the size of the data point.

**S2 Fig.**
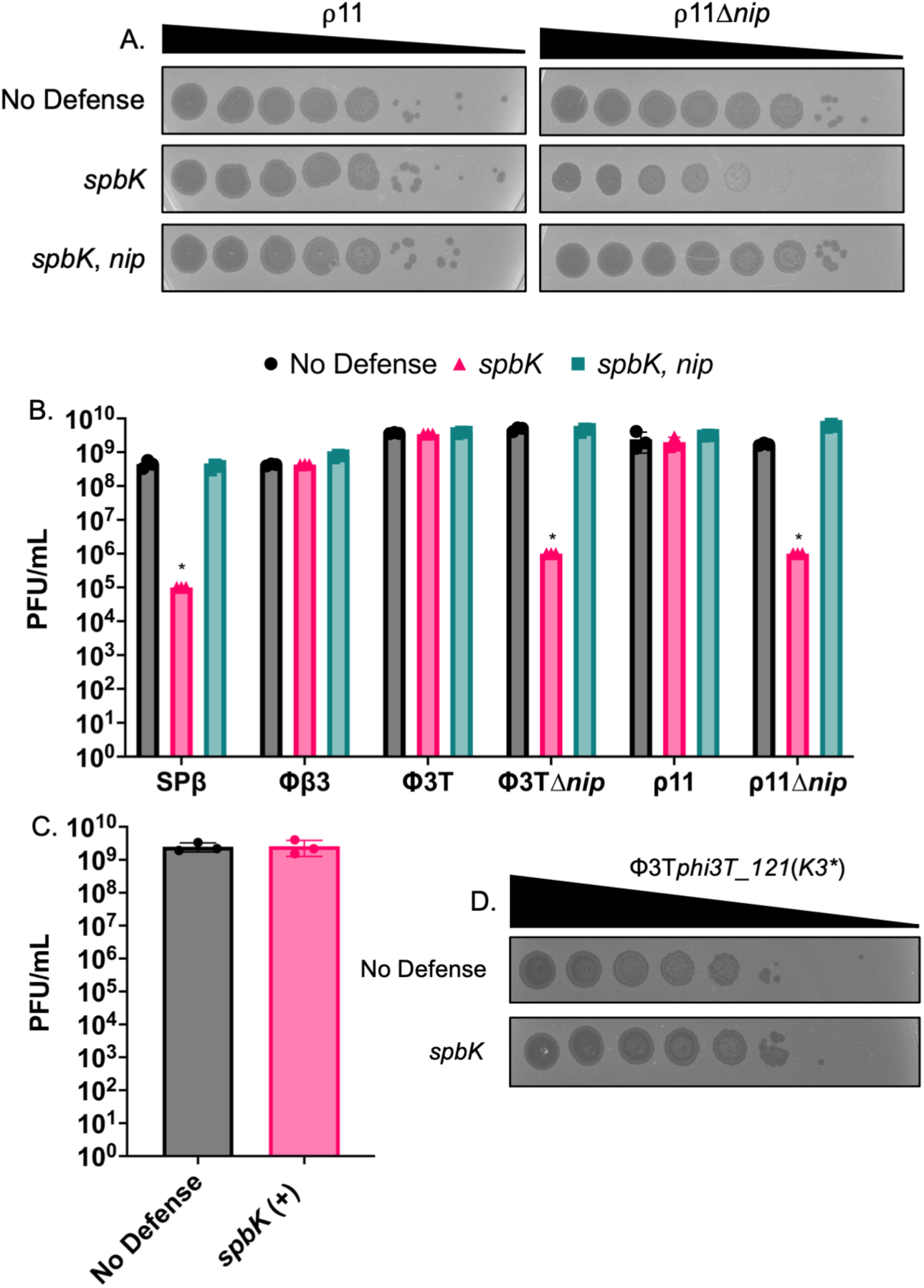
*nip* is necessary and sufficient to prevent SpbK-mediated phage defense in ρ11, SPβ, and Φ3T. **A)** *nip* is necessary and sufficient to enable ρ11 evasion of SpbK-mediated anti-phage defense. Ten-fold dilutions of ρ11 (left) or ρ11Δ*nip* (right) were spotted onto isogenic strains without phage defense (CU1050, top), expressing *spbK* (CMJ534, middle), or co-expressing *spbK* and *nip* (CLL356; bottom). Large zones of clearing indicate cell lysis while small zones of clearing are indicative of productive phage infection. **B)** *nip* prevents SpbK-mediated phage defense. Strains lacking anti-phage defense (black bars), expressing *spbK* (pink bars), or co-expressing *spbK* and *nip* (turquoise bars) were infected with phage indicated at the x-axis and efficiency of plaquing was measured. Asterisks indicate limit of detection. Data are from three biological replicates. **C)** Φ3T*phi3T_121*(*K3STOP*) does not exhibit a defect in plaquing efficiency. Strains lacking anti-phage defense (black bar) or expressing *spbK* (pink bar) were infected with Φ3T*phi3T_121*(*K3STOP*). Error bars represent standard deviation and are not always visible. **D)** Ten-fold dilutions of Φ3T*phi3T_121*(*K3STOP*) were spotted onto isogenic strains lacking anti-phage defense (CU1050, top row) or expressing *spbK* (CMJ534, bottom row). Large zones of clearing indicate cell lysis while small zones of clearing are indicative of productive phage infection.

**S1 Data.** Underlying raw data for experiments presented in the graphs. The excel spreadsheet contains the underlying data for the experiments presented in each of the graphs.

**S3 Fig.**
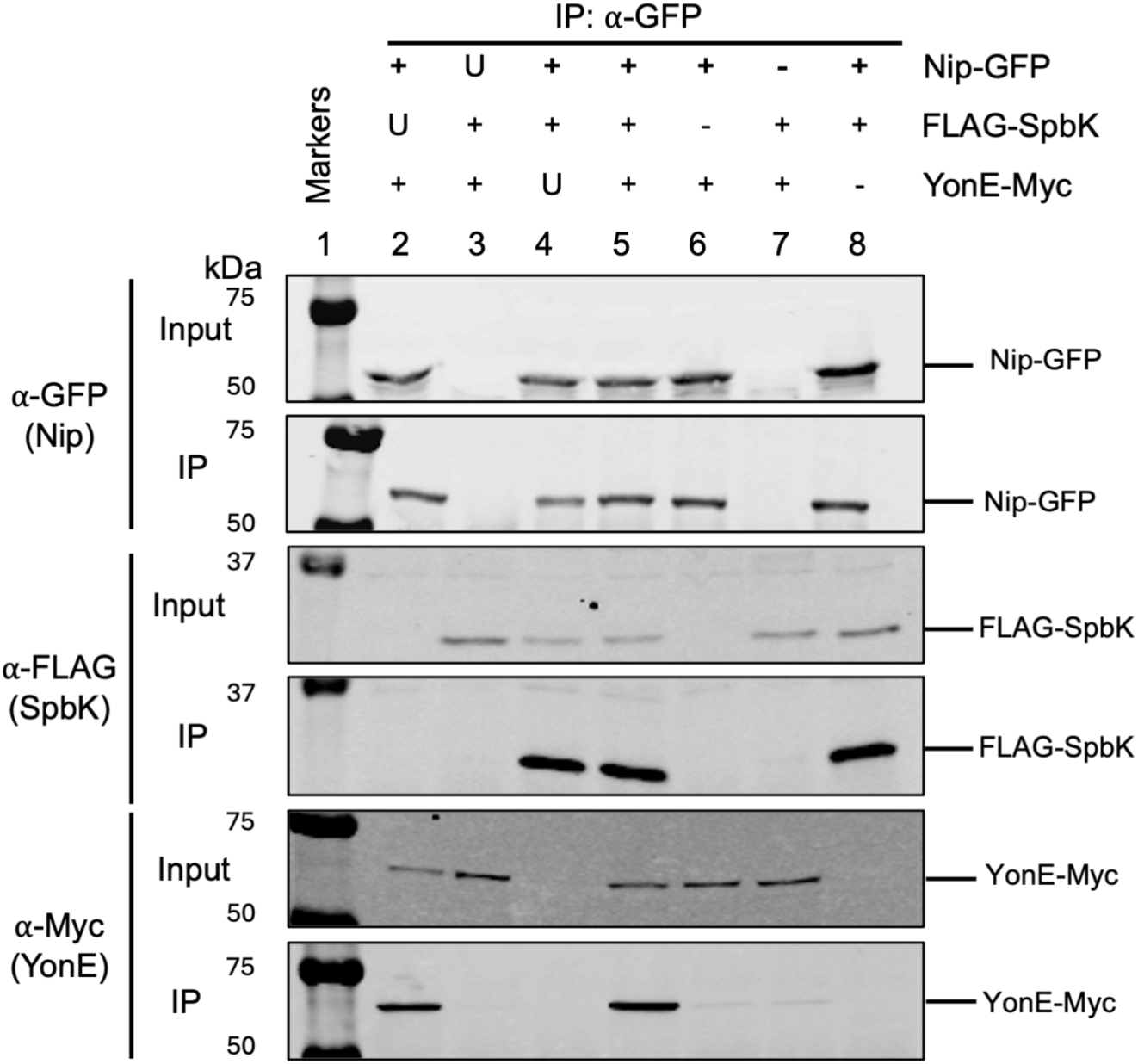
**Nip, SpbK, and YonE form a tripartite complex: Input and immunoprecipitation samples.** Data for the immunoprecipitation experiments are the same as in Fig 7 and are shown here for comparison. Input samples are shown in the panel above each of the corresponding immunoprecipitations (IP). Indicated proteins were expressed in cells without ICE*Bs1* or SPβ. First and second rows: Western blots probed with α-GFP polyclonal antibodies on input samples and immunoprecipitates, respectively. Third and fourth rows: Western blot probed with α-FLAG monoclonal antibodies on input samples and immunoprecipitates, respectively. Fifth and sixth rows: Western blot probed with α-c-Myc monoclonal antibodies on input samples and immunoprecipitates, respectively. Lane 1 contains molecular weight markers from the Odyssey One-Color Protein Molecular Weight Marker Ladder (LI-COR). Lysates and IPs were from strains: CLL633 (lane 2), CLL642 (lane 3), and CLL704 (lane 4), CLL498 (lane 5), CLL497 (lane 6), CLL373 (lane 7), and CLL382 (lane 8). Data shown are representative of three biological replicates. U, untagged protein expressed; +, tagged protein expressed; -, protein not expressed.

